# Extracellular polymeric substance degradation shapes microbial community diversity

**DOI:** 10.1101/2025.01.22.634264

**Authors:** Sammy Pontrelli, Kian Bigovic Villi, Andreas Sichert, Julian Trouillon, Adriano Rutz, Zachary Landry, Simon H. Rüdisser, Roman Stocker, Uwe Sauer

## Abstract

Competition for and exchange of nutrients play crucial roles in shaping microbial community function and dynamics. Although cross-feeding of small metabolites is known to drive carbon exchange among species, the importance of self-produced extracellular polymeric substances (EPS)— which include proteins, polysaccharides, DNA, and humic-like substances— remains less understood. Utilizing chitin-degrading microbial isolates and natural seawater communities, we found that 4–7% of the carbon from chitin degradation is converted into EPS, accounting for nearly a quarter of the exuded carbon. Different sources of EPS were found to select for distinct microbial communities. Through enzyme assays and untargeted metabolomics, we demonstrated that secreted enzymes degrade EPS in multiple steps that influence community diversity: larger oligomers are initially utilized by specialized degraders, while the subsequent breakdown into smaller oligomers, monosaccharides, and amino acids supports non-specialized consumers. These findings highlight the role of EPS as a significant carbon source exchanged between microbes, fueling metabolically diverse populations and enriching our understanding of carbon-mediated microbial interactions.

## Introduction

Microbial communities are vital for biogeochemical cycles and the health of humans, animals and plants, as their structure and composition influence ecosystem functions^1,2^. Research investigating the factors that shape community structure has revealed that specialized species, which can utilize complex carbon sources, produce metabolites that contribute to the formation of cross-feeding networks^3–5^. These species supply carbon to less specialized community members, thereby shaping temporal population dynamics, and defining community composition^3,5–7^. Traditionally, research on metabolite cross-feeding has focused on low-molecular-weight metabolites, such as monosaccharides, organic acids, amino acids, and vitamins^8–10^. These metabolites are fundamental building blocks of life, often secreted in large quantities by microbes in various environments, and have been found to directly drive certain species interactions^3,8,9,11^. Consequently, small metabolites have become central to mass spectrometry-based metabolomics studies of cross-feeding.

Many species produce extracellular polymeric substances (EPS) that can serve as potential nutrients shared among species^12,13^, but these compounds are not directly accessible through metabolomics methods. Due to their size and complexity, high-molecular-weight EPS may be overlooked in classical cross-feeding studies. EPS encompasses a variety of components, including polysaccharides, nucleic acids, proteins, and humic-like substances, each secreted for specific functions^13^. Their ubiquity across diverse ecosystems make them readily available as carbon substrates. For instance, biofilms are primarily defined by polysaccharides with varying chemical features^14–16^; secreted proteins catalyze degradation of external polysaccharides^17,18^; and humic-like substances naturally form during microbial degradation of organic matter^19^. The widespread availability of EPS may drive an evolutionary selection for species capable of utilizing its components, suggesting that EPS could be a universal factor in shaping microbial community structures. Several studies have provided evidence that EPS can select for specific community members and alter community structure. For instance, marine microbial communities cultured on concentrated, bacterial-derived EPS tends to favor distinct species^20^. Similarly, diatom-derived EPS, extracted through various methods that yield concentrates with differing chemical compositions, can alter species abundances when supplemented into marine sediment microbial communities^21^. Consequently, EPS and its specific components play a critical role in shaping population dynamics. However, the chemical fractions that mediate carbon exchange, their relative contribution to carbon flux, and their overall impact on community structure remain poorly understood^13^.

This study investigates how EPS mediates carbon exchange among microbes, influencing population dynamics within a chitin-degrading microbial community. Chitin, an insoluble polysaccharide found in crustacean and copepod shells, is one of the most abundant polysaccharides in the ocean and plays a critical role in the global carbon cycle^22^. Previous research has demonstrated that cellular metabolites and chitin breakdown products released by specialized chitin degraders have a cascading influence on the population dynamics of subsequently colonizing species^3,23,24^. However, the role of EPS in this context has not been explored. Here we reveal that high-molecular-weight compounds constitute a major portion of exuded carbon, and that different sources of EPS distinctly shape community composition. While only specialized microbes can degrade EPS, the extracellular enzymes they release break EPS down into smaller entities that become accessible to non-specialized species, thereby fostering community diversity. These findings enhance our understanding of metabolic cross-feeding by illustrating how high-molecular-weight compounds drive carbon exchange and influence community dynamics.

### EPS yields and compositions during chitin degradation

To assess the potential role of EPS in carbon flux between degraders and non-degraders, we quantified the proportion of chitin-derived carbon allocated to EPS (larger than 3 kDa), low-molecular-weight (LMW) compounds (below 3 kDa), and biomass in the model chitin-degrading bacterium *Vib*1A01^11,25,26^. Upon reaching early stationary phase on chitin as the sole carbon source, we collected biomass, separated EPS from LMW fractions using a 3 kDa filter, and measured the carbon content in each fraction. Of the 2 g/L chitin that were completely consumed (data not shown), 33-36% of carbon were recovered in biomass, 4.1-4.5% in EPS, and 11-14% in LMW compounds, while the remaining unaccounted 45-52% were deemed lost as CO_2_ in respiration. EPS secretion in two other chitin-degrading isolates was similar **(Fig. 1A)**. When chitin cultures were inoculated with non-sterilized seawater, we observed comparable EPS production of 4-7% of the total chitin carbon, independent of variations in community composition as determined through 16S sequencing (**Fig. 1B**, Bray-Curtis dissimilarity: 0.78 and 0.69 for communities 1 vs. 2 and 3, and 0.31 for 2 vs. 3). Since EPS consistently accounts for about a quarter of exuded carbon from bacterial isolates and communities, it has considerable potential to contribute to carbon flux within chitin degrading communities.

**Figure 1:**
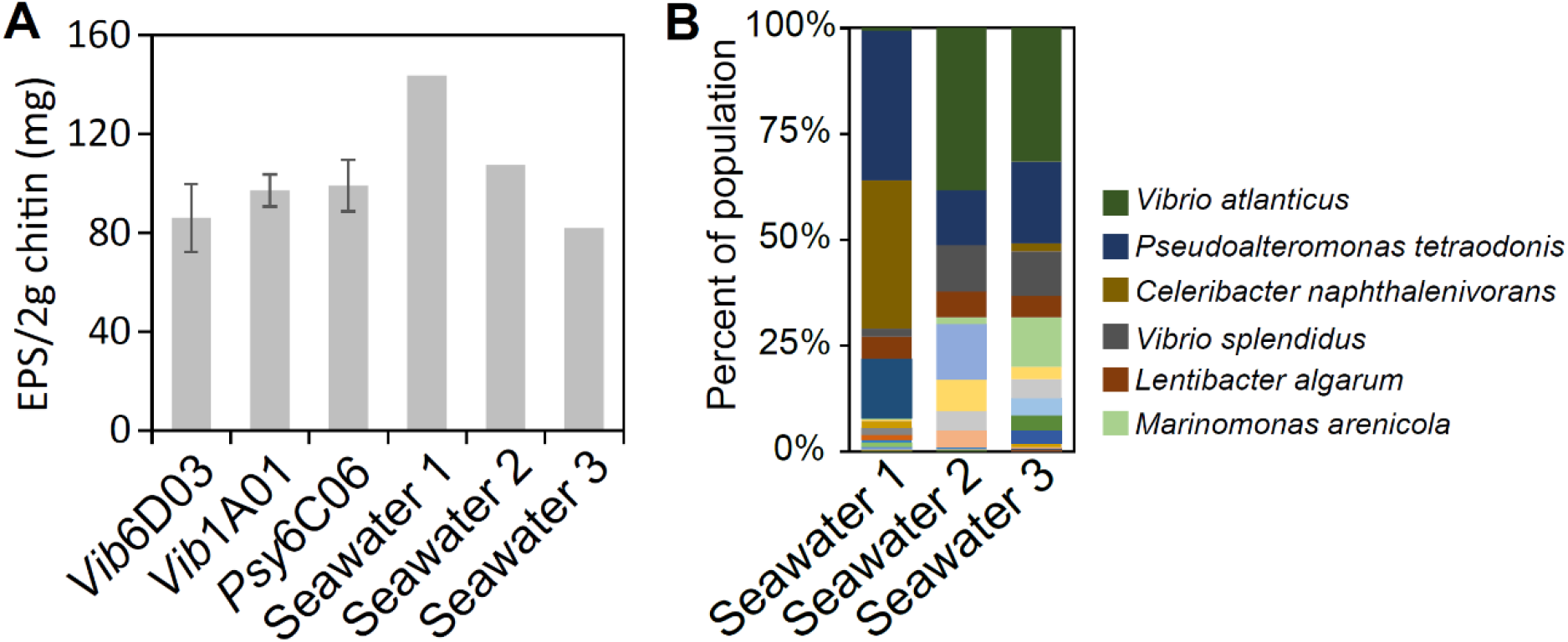
EPS yields from chitin. A) EPS concentrations in early stationary phase after growth on 2 g/L of chitin as the sole carbon source. The three species are known chitin-degraders and the three seawater inocula. Error bars represent the standard deviation of the mean from three biological replicates. B) 16S sequences of the three seawater communities at the time of EPS harvesting, showing the average top 6 most abundant species.

Next, we investigated the molecular composition of the six EPS fractions because EPS are a heterogeneous mix of macromolecules^13^. In degrader monocultures, proteins, which are naturally secreted for chitin degradation, comprised 13–23% (g/g) of the EPS. In seawater communities, proteins accounted for less than 3% or were undetectable **(Fig. 2A)**, suggesting that proteins may have been consumed by non-degrading microbes. Polysaccharides, which play critical roles in cell adhesion, bacterial aggregation, and biofilm formation, were subjected to acid hydrolysis, and the resulting monosaccharides were quantified using liquid chromatography-mass spectrometry (**Supplemental Table 1)**. The polysaccharide content varied widely, representing 9–36% of the total carbon in the six EPS samples **(Fig. 2B)**. Glucose emerged as the most abundant monosaccharide in five of the samples, averaging 55% of the monomers by weight. Overall, proteins and hydrolysable polysaccharides constituted 22–59% (g/g) of the exuded EPS.

**Figure 2:**
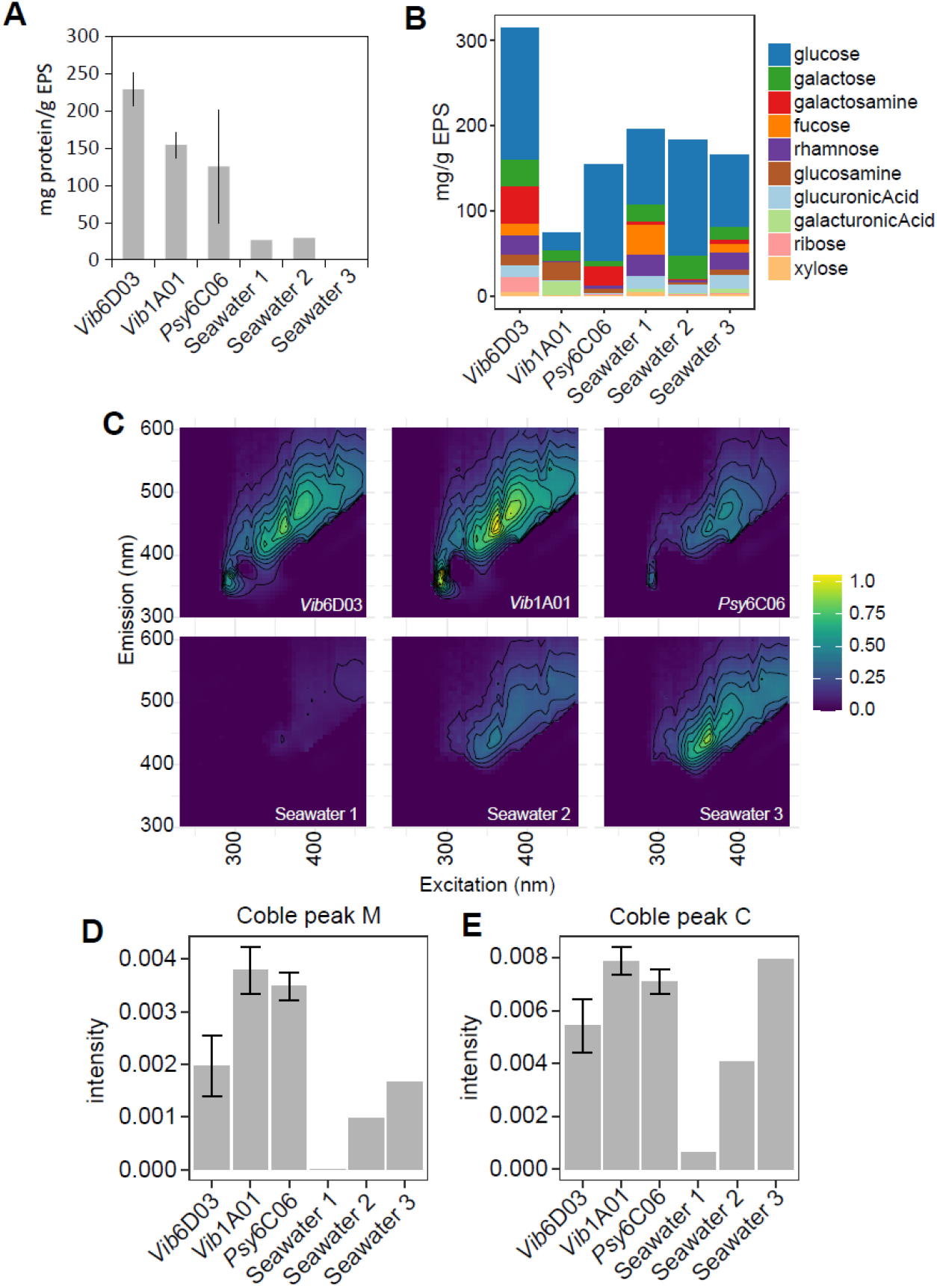
Composition of EPS. A) Mass fraction of proteins in EPS from different sources. B) Mass fraction of monosaccharides in hydrolysable polysaccharides in EPS from different sources **(Supplemental Table 1)**. C) Excitation emission matrices of each EPS at 1 g/L. D) Coble peak M (Ex/Em 290-310/370-410) and peak C (Ex/Em 320-360/420-460) in the excitation emission matrices. Error bars represent standard deviation of the mean of three biological replicates.

High-molecular-weight humic-like substances, which form naturally during bacterial degradation of organic matter^27^, can be important components of EPS that may also support specific microbes^28^. Their complex structures complicate quantification; however, fluorescent signatures, such as the Coble M peak (Ex 290-310 nm, Em 370-410 nm) for marine and Coble C peak (Ex 320-360 nm, Em 420-460 nm) for visible humic-like substances, can revealed their presence ^29^. We obtained excitation-emission matrices and detected both C and M peaks in the EPS of all three degraders **(Fig. 2CDE)**. In two of the three seawater samples, however, the C and M peaks were much lower than in the monocultures. These variations in humic-like substances, along with differences in protein and polysaccharide content, illustrate diverse EPS composition across different sources of EPS.

### Influence of EPS composition of microbial community structure

Given the variable EPS composition, we hypothesized that their unique components would select for distinct microbial communities of heterotrophic bacteria. To test this hypothesis, we cultured microbes naturally found in seawater using EPS as the sole carbon source and analyzed the community composition through 16S sequencing. To prevent the formation of entirely distinct community assemblages from the same dilute seawater inoculum (as shown in Figure 1B), we initially prepared the inoculum by inoculating a minimal medium supplemented with five different carbon sources with seawater. This approach generated a dense and diverse culture, which was then used to seed the various EPS media. As a result, any observed changes in community composition can be attributed specifically to the different EPS substrates rather than to random inoculum variations.

After 48 hrs, robust growth was observed on all EPS substrates **(Fig. 3A)**. While communities shared some abundant taxa, each assembled with distinct population structures, uniquely enriched with specific degraders compared to the inoculum **(Fig. 3BC)**. Notably, *Vibrio gigantis*, a marine polysaccharide degrader^30^, increased 7-22 fold in five out of the six EPS conditions when compared to the inoculum. In the sixth condition, *Alteromonas stellipolaris*, a representative of the *Alteromonas* genus known for its ability to degrade algal polysaccharides^31^ and polycyclic aromatic hydrocarbons^32^, increased nearly 9 fold. Moreover, six *Pseudoalteromonas* species, known for degrading marine polysaccharides and proteins^33,34^, increased up to 5 fold exclusively in *Psy*6C06 EPS, demonstrating EPS specificity in selecting distinct EPS-degrading microbes .To further demonstrate that EPS degradation is a specialized trait, we screened the growth of 18 bacteria from a model chitin-degrading community^3^ on the EPS of *Vib*1A01 as the sole carbon source **(Table 1)**. While most species displayed some growth, *Alt*A3R04 and *Mar*F3R11 grew to the highest OD_600_ **(Fig. 3E)**. Overall, these results show that different EPS types select for specific degraders, which may subsequently influence community structure.

**Figure 3:**
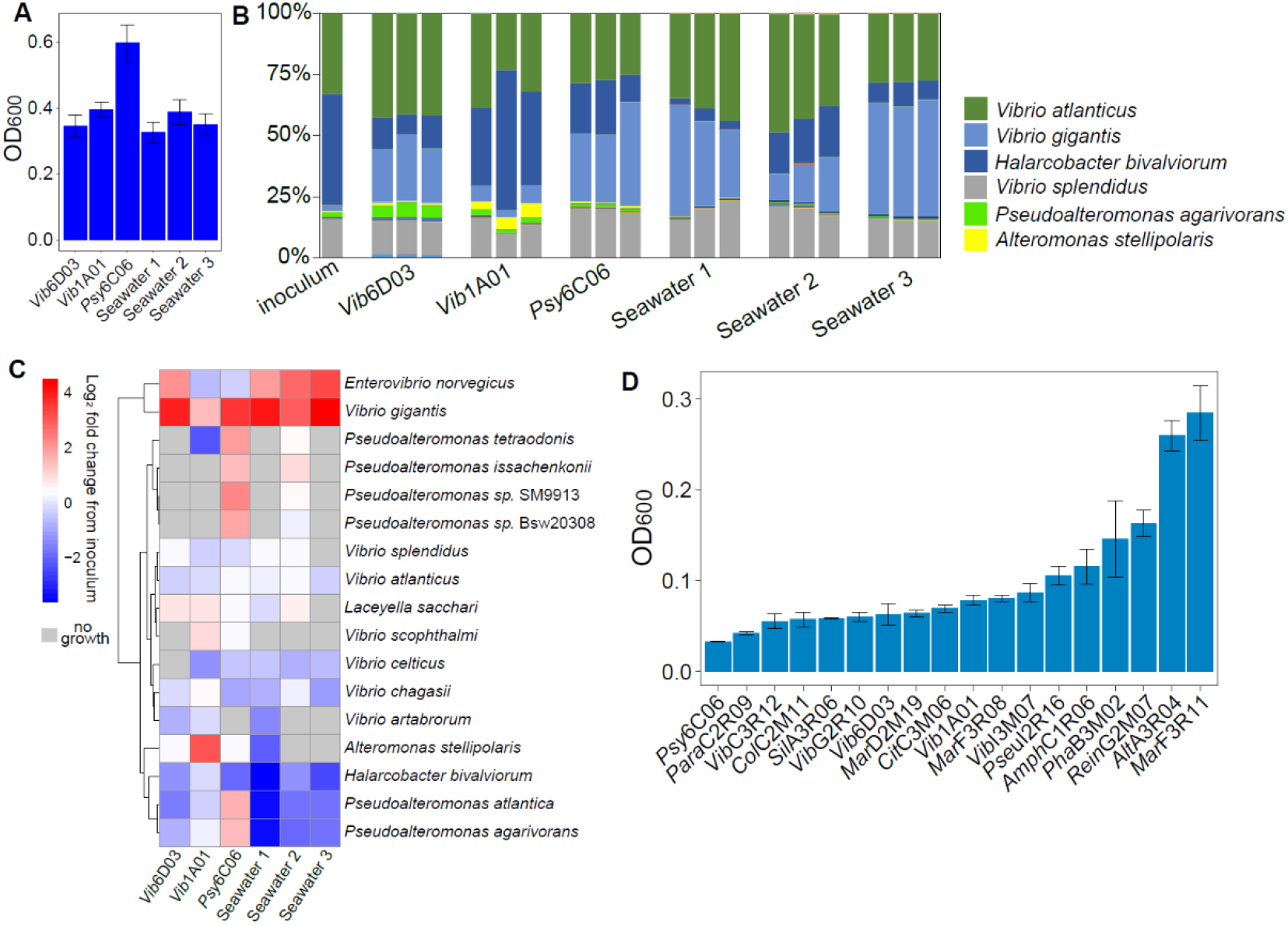
Growth selection by EPS as a carbon source. A) Optical density 48 h after inoculation with seawater on 1 g/L EPS as a sole carbon source, derived from the indicated chitin degrading cultures. B) 16S sequencing of seawater communities after growth on each EPS medium. C) Heatmap showing Log^2^ fold change in abundance of species in the EPS communities compared to the inoculum, where species not found in the EPS communities shown in gray. Species are clustered using Ward’s method. D) Growth of individual bacterial isolates on 1 g/L of EPS from *Vib*1A01 as a sole carbon source at 72 hrs. All error bars represent standard deviation of three biological replicates.

**Table 1:** Species used in this study and NCBI accession numbers for genomic sequences. The functional guild describes whether species have the ability to degrade chitin and grow on it as a sole carbon source.

| Species | Taxonomic ID | Functional Guild | NCBI BioProject | NCBI BioSample |
| --- | --- | --- | --- | --- |
| AltA3R04 | *Alteromonas sp.* | Non-degrader | PRJNA478695 | SAMN19350919 |
| AmphC1R06 | *Amphritea sp.* | Non-degrader | PRJNA478695 | SAMN19350932 |
| CitC3M06 | *Citricella sp.* | Non-degrader | PRJNA478695 | SAMN19350936 |
| ColC2M11 | *Colwellia psychrerythraea* | Non-degrader | PRJNA478695 | SAMN19350933 |
| MarD2M19 | *Marinobacter sp.* | Non-degrader | PRJNA478695 | SAMN19350941 |
| MarF3R08 | *Marinobacter sp.* | Non-degrader | PRJNA478695 | SAMN19350954 |
| MarF3R11 | *Marinobacter sp.* | Non-degrader | PRJNA478695 | SAMN09522136 |
| ParaC2R09 | *Paracoccus kamogawaensis* | Non-degrader | PRJNA478695 | SAMN19350935 |
| PhaB3M02 | *Phaeobacter sp.* | Non-degrader | PRJNA478695 | SAMN09522130 |
| Pseul2R16 | Pseudoalteromonadaceae | Non-degrader | PRJNA478695 | SAMN19350960 |
| Psy6C06 | *Psychromonas sp.* | Degrader | PRJNA414740 | SAMN08130274 |
| ReinG2M07 | *Reinekea sp.* | Non-degrader | PRJNA478695 | SAMN19350955 |
| SilA3R06 | *Silicibacter sp.* | Non-degrader | PRJNA478695 | SAMN19350920 |
| Vib1A01 | *Vibrio splendidus* | Degrader | PRJNA414740 | SAMN07809270 |
| Vib6D03 | *Vibrio penaeicida* | Degrader | PRJNA414740 | SAMN08130373 |
| VibC3R12 | *Vibrio sp.* | Non-degrader | PRJNA478695 | SAMN09522132 |
| VibG2R10 | *Vibrio sp.* | Degrader | PRJNA478695 | SAMN19350956 |
| VibI3M07 | *Vibrio sp.* | Degrader | PRJNA478695 | SAMN19350961 |

### Stepwise enzymatic degradation of EPS

While EPS degradation appears to rely primarily on specialized microbes, the diversity and presence of certain species across all EPS conditions **(Fig. 3BC)** suggest that non-specialized, non-degrading microbes are also supported. One possibility is that these species obtain nutrients by exploiting the extracellular breakdown of EPS into monomer or oligomers without themselves releasing enzymes^3,35,36^. To assess the range of EPS degradation products that may fuel these species, we purified EPS from the three chitin degraders after they were grown on colloidal chitin and subsequently used it as a sole carbon source for a seawater community. Once the cultures reached early stationary phase, we removed cells by centrifugation and concentrated secreted proteins along with any remaining EPS. This concentrated enzyme mixture was then incubated *in vitro* with fresh EPS, and aliquots were withdrawn over a 24 h period to monitor the formation of degradation products using untargeted liquid chromatography quadrupole time of flight mass spectrometry (LC-QTOF-MS) mass spectrometry-based metabolomics.

A total of 2820 ions displayed one of three temporal patterns in at least one of the enzyme digests **(Fig. 4A)**. Firstly, 1226 ions increased over time, indicating the accumulation of digestion products. Secondly, 492 ions showed a decreasing trend, with most of these being larger molecular weight ions, suggesting their breakdown during the incubation. Thirdly, 1279 ions displayed bell shape curves, suggesting an initial formation of oligomers followed by subsequent degradation. Upon analyzing the size distribution of ions in each temporal category **(Fig. 4B)**, we found that both the decreasing ions and those with bell-shaped curves were predominantly composed of larger ions (1600-1800 *m/z*). Conversely, ions increasing over time were predominantly small ions (200-400 *m/z*). These results show that extracellular enzymes initially break down EPS into large oligomers, which are then further degraded, presumably due to collaborative action of multiple enzymes. To determine whether EPS degradation products are degraded into unique oligomers or more common components, we analyzed the ions that increased over time. Among the ions with *m/z* values above 1400, only 15% were present in multiple digests, while over half of smaller ions (50–200 *m/z*) were found across various digests **(Supplemental Dataset 1)**. Together, this evidence suggests that enzymes initially degrade EPS into unique oligomers that select for specific species, whereas further degradation produces common monomers and oligomers. These smaller molecules likely require less metabolic specialization, enabling less specialized species to grow by exploiting extracellular EPS degradation initiated by degraders **(Fig. 4C)**.

**Figure 4:**
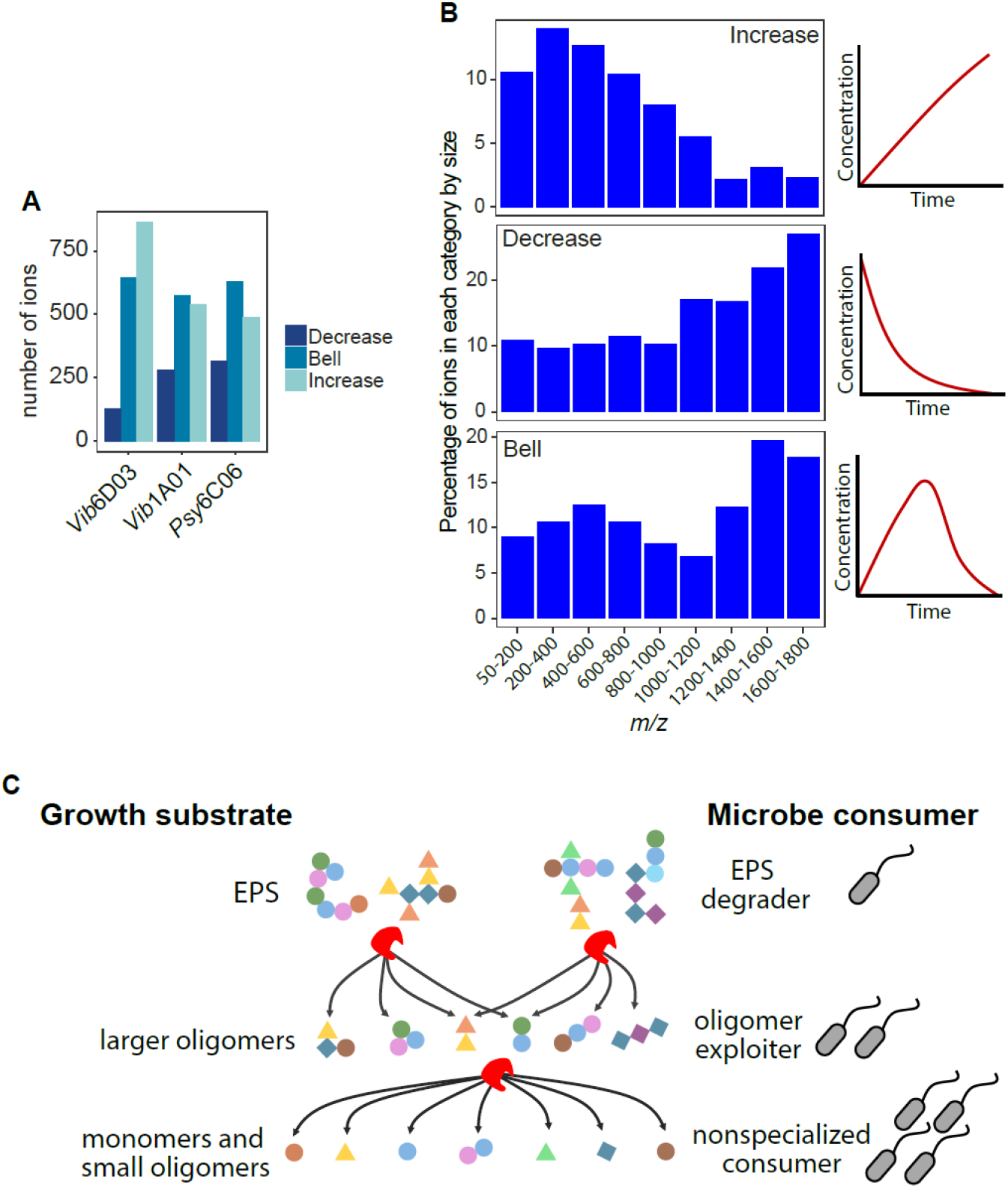
Size and temporal dynamics of EPS digestion. A) Number of ions in each enzyme digest falling in the decreased, bell-shaped, or increased categories. B) Percentage of all detected ions in different size ranges that fall into the three temporal categories. C) Graphical illustration of how EPS is enzymatically degraded in multiple steps into smaller fragments, fueling the growth of progressively less specialized species: Specialized EPS degraders break down EPS into larger oligomers for oligomer exploiters; further degradation into monomers and small oligomers supports nonspecialized consumers.

To identify specific degradation products, we putatively annotated 279 of the 2,820 detected ions based on their exact masses using the BioCyc database^37^ **(Supplemental Dataset 2)**, and used CANOPUS^38^ to classify them into more detailed compound classes ions for which MS^2^ spectra were acquired **(Supplemental Dataset 3)**. Consistent with proteins accounting for 13–23% of EPS **(Fig. 3A)**, the concentrations of the 17 detectable amino acids increased by at least two-fold in one or more digests, with 7 amino acids showing increases in all digests **(Fig. 5A)**. This indicates a common release of amino acids in all digests. We further annotated peptides by examining the compound classes related to their fragmentation spectra or by comparing the exact masses of detected ions to those of peptides containing four or fewer amino acids. These peptides exhibited three distinct temporal patterns: increasing, decreasing, or bell-shaped **(Fig. 5B)**. This supports the conclusion that EPS proteins are sequentially degraded into peptides, which are ultimately broken down into amino acids that provide nutrients for non-specialist microbes. Additionally, we observed accumulation of putatively annotated acetyl-hexosamine and aminohexose dimers in the digests **(Fig. 5C)**, along with acetyl-hexosamine and O-glycosyl compound classes **(Fig. 5D)**, likely arising from the degradation of EPS polysaccharides. Consistently, NMR analysis of undigested *Vib*1A01 EPS confirmed strong peaks for acetylated sugars and carbohydrates **(Fig. 5E)**, as well as for proteins. Akin to the amino acids, hexosamines and acetyl-hexosamines can support the growth of both specialized degraders and less-specialized microbes once the polysaccharides are degraded.

**Figure 5:**
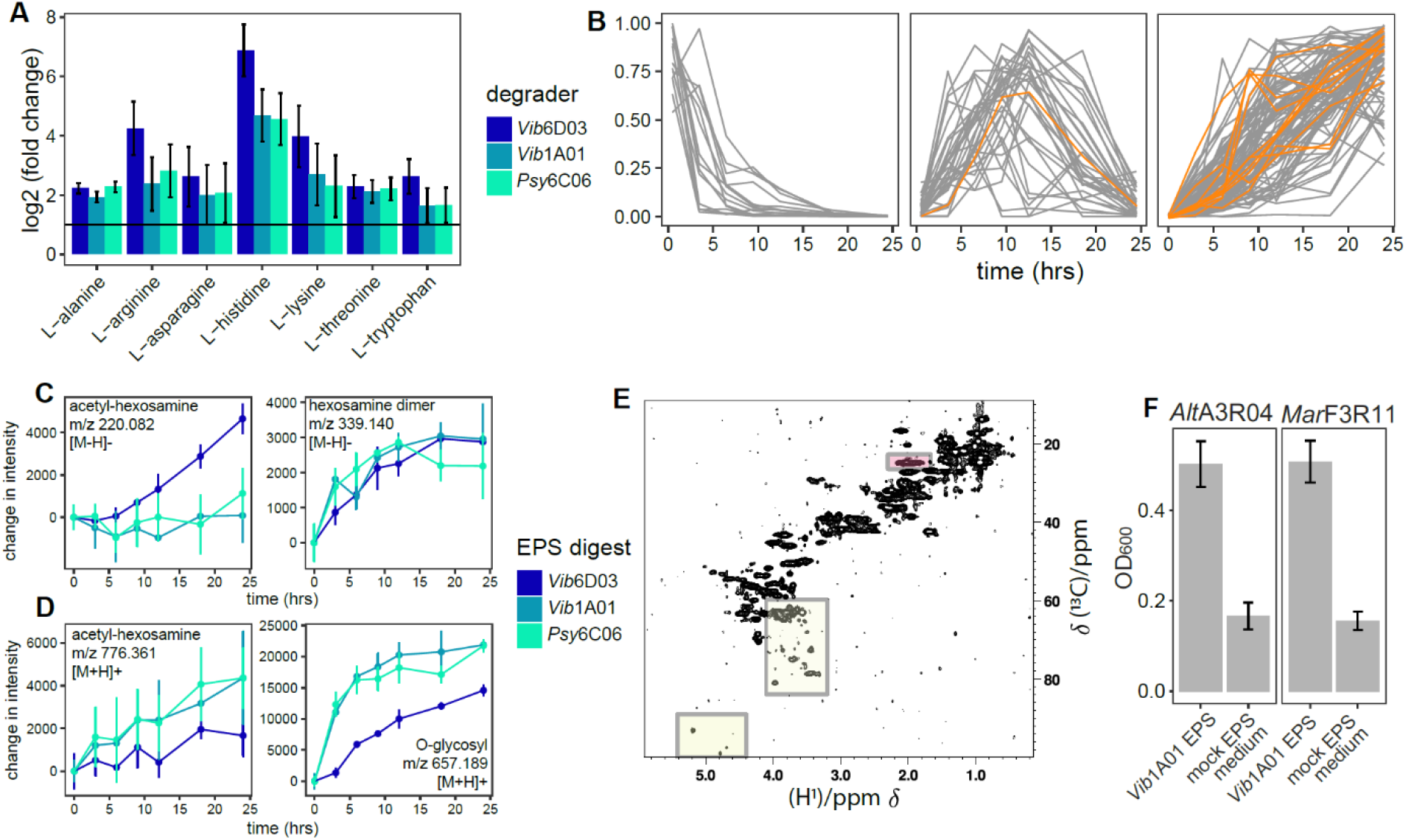
Release of amino acids and monosaccharides from EPS. A) Log2 fold change of amino acid concentrations comparing 0 and 24 hrs of the enzyme digests. B) Temporal trajectories of annotated peptides with 4 or fewer amino acids in the enzyme digest of *Vib*6D03, with ions in orange annotated as peptides or amino acids based on MS^2^-based CANOPUS predictions. Intensity change of compounds in enzyme digests relative to time 0 for C) acetyl-hexosamine and hexosamine dimers, annotated by exact mass, and D) ions corresponding to acetyl-hexosamine and O-glycosyl compound classes, annotated by CANOPUS. E) ^1^H-^13^C HSQC NMR spectrum of EPS. The spectral regions of the carbohydrate resonances are highlighted by yellow boxes. The signal at the chemical shift which is characteristic for an N-acetyl group is highlighted in red. The signals outside of these regions indicate the presence of amino-acids, peptides or small proteins. F) Growth of EPS degrading microbes on 1 g/L of *Vib*1A01 enzyme digest and mock EPS medium containing proteins and monosaccharides to mimic those found in *Vib*1A01 EPS. Error bars represent standard deviation of the mean of three biological replicates.

The metabolomics data show that proteins and polysaccharides could potentially feed EPS-degrading communities. To investigate whether these two polymers are primarily responsible for observed growth, or if other EPS components also contribute, we designed a medium that mimics 1 g/L of *Vib*1A01 EPS. This medium contains 150 mg/ml of protein and the approximate composition of all monosaccharides present at concentrations greater than 1 µM from the polysaccharide fraction **(Supplemental Table 1)**. The EPS degrading isolates *Alt*A3R04 and *Mar*F3R11 reached 30–33% of the final density in the formulated medium compared to the actual EPS from *Vib*1A01 **(Fig. 5F)**. Our mimicry medium likely underestimates the role of proteins and especially polysaccharides in promoting growth, primarily due to incomplete monosaccharide identification and limitations in distinguishing between unmodified and functionalized (e.g., acylated) polysaccharides with higher carbon content. Nevertheless, our results demonstrate that proteins, polysaccharides, and unidentified EPS components all serve as important cross-fed nutrient sources in chitin-degrading communities.

## Discussion

In this work, we demonstrate that EPS plays a significant role in carbon exchange between chitin-degraders and non-degrading consumers, both in terms of quantity of carbon available for growth and its influence on community dynamics. Our findings reveal that 4–7% of the carbon released by degraders is converted into EPS, which, in the case of the model chitin-degrader *Vib*1A01^3,11,25^, accounts for approximately a quarter of the total carbon secreted. Different EPS sources resulted in distinct microbial assemblages when fed to seawater communities and promoted different degraders with varying abundances. This mirrors previous studies that have shown that enzymatic degradation of chitin into N-acetyl glucosamine oligomers of varying chain lengths influences the selection of species capable of consuming these different oligomers^3^. Given that EPS exhibits greater complexity than homogenous polymers composed of a single monomer type, such as chitin, its degradation is likely to yield a wider diversity of oligomers, which in turn supports more specialized oligomer consumers.

While small cross-fed metabolites are known to mediate carbon exchange within diverse microbial communities^3,9,11^, the presence of overlapping strains across various EPS-grown communities suggests that also EPS contributes to community diversification. Our findings demonstrate that when EPS is broken down into oligomers, it undergoes further degradation into small monomers in a step-wise fashion, producing common and easily metabolizable substrates that can sustain non-specialized consumers. These observations aign with previous studies of microbial community assembly, which show that polysaccharide-degrading communities exhibit modular dynamics^23,39^. Initially, distinct microbial populations colonize specific polysaccharides, but, over time, these communities converge to support generalist species. By showing that microbial EPS secretion and its breakdown into simpler carbon sources by other species drive colonization patterns, our research underscores the importance of incorporating EPS in studies that link carbon exchange to microbial community structure.

By employing analytical tools to quantify EPS components, along with enzyme assays and metabolomics to identify degradation products, we show that EPS proteins and polysaccharides substantially support microbial community growth. Furthermore, we show that other EPS components not identified in this study likely contribute to carbon exchange. These may include humic-like substances, which possess undefined structures that complicate their quantification. Additionally, non-hydrolysable polysaccharides, such as acylated deoxy sugars - which cannot be broken down chemically and thus are difficult to quantify - may constitute a significant fraction of EPS and contribute to marine dissolved organic matter^40^. Current research efforts are aimed at identifying the composition of these compounds and elucidating their roles in marine carbon cycling. Our findings emphasize the importance of understanding this composition not only in the context of carbon cycling but also regarding its impact on microbial community structure and function.

## Methods

### Materials and chemicals

All chemicals, unless otherwise specified, were purchased from Sigma-Aldrich. The media used include Marine Broth 2216 (Thermo Fisher Scientific, Difco, no. 279100) and MBL minimal medium.

MBL medium consists of 1 mM phosphate dibasic, 1 mM sodium sulfate, and 50 mM TES (pH 8.2), along with three additional diluted stock solutions. Fourfold concentrated seawater salts (NaCl, 80 g/L; MgCl_2_*6H_2_O, 12 g/L; CaCl_2_*2H_2_O, 0.6 g/L; KCl, 2 g/L). 1000-fold concentrated trace minerals (FeSO_4_*7H_2_O, 2.1 g/L; H_3_BO_3_, 30 mg/L; MnCl_2_*4H_2_O, 100 mg/L; CoCl_2_*6H_2_O, 190 mg/L; NiCl_2_*6H_2_O, 24 mg/L; CuCl_2_*2H_2_O, 2 mg/L; ZnSO_4_*7H_2_O, 144 mg/L; Na_2_MoO_4_*2H_2_O, 36 mg/L; NaVO_3_, 25 mg/L; NaWO_4_*2H_2_O, 25 mg/L; Na_2_SeO_3_*5H_2_O, 6 mg/L, dissolved in 20 mM HCl). 1000-fold concentrated vitamins (riboflavin, 100 mg/L; d-biotin, 30 mg/L; thiamine hydrochloride, 100 mg/L; l-ascorbic acid, 100 mg/L; Ca d-pantothenate, 100 mg/L; folate, 100 mg/L; nicotinate, 100 mg/L; 4-aminobenzoic acid, 100 mg/L; pyridoxine HCl, 100 mg/L; lipoic acid, 100 mg/L; nicotinamide adenine dinucleotide (NAD), 100 mg/L; thiamine pyrophosphate, 100 mg/L; cyanocobalamin, 10 mg/L, dissolved in 10 mM MOPS, pH 7.2).

### Preparation of colloidal chitin

Ten grams of powdered chitin (Sigma-Aldrich, C7170) was dissolved in 100 ml of concentrated phosphoric acid (85% (w/v)) and incubated at 4°C for 48 hrs. Approximately 500 ml of deionized water was added to this mixture and shaken vigorously until all the chitin precipitated. The precipitate was filtered using regenerated cellulose paper (MACHEREY-NAGEL, MN615). The chitin precipitate was then placed in cellulose dialysis tubing (approximately 13 kDa, Sigma-Aldrich D9652-100FT) and dialyzed with fresh deionized water daily for three days to remove residual phosphoric acid and oligomers. After dialysis, the pH was adjusted to 7 with 1M NaOH and homogenized using a Bosch SilentMixx Pro blender. The colloidal chitin was sterilized by autoclaving.

### Bacterial strains and culturing

All bacterial species were stored in glycerol stocks at −80°C. Before use, they were streaked onto Marine Broth 2216 plates with 1.5% (w/v) agar (BD, no. 214010) and incubated at room temperature until colonies formed. Overnight precultures were prepared by inoculating a single colony into 2 ml of Marine Broth 2216 and incubating in a 27°C shaker overnight. In the case of *Psy*6C06, which grows to a low optical density in MB 2216 (OD_600_ ∼0.2), the 2 mL preculture was used to inoculate a second overnight preculture of 15 mL.

Unless otherwise noted, growth experiments were performed as follows. The medium used was MBL. Cells were inoculated at a density of 1×10_7_ cells/mL. When chitin was used as a carbon source, it was supplied at 2 g/L to 15 mL cultures in 100 mL shake flasks. In all other cases, culture volumes were 200 µL in a 96 well plate, where all wells on the rim of the plate were filled with water to minimize volume loss to evaporation. When EPS is used as a carbon source it was added at 1g/l.

Growth was monitored by measuring OD_600_. For chitin cultures, 200 μL of culture was harvested for measurement in a Tecan Sunrise plate reader. To prevent suspended chitin from interfering with OD_600_ measurements, the sample was centrifuged for 30 seconds at 1000 rcf, and the optical density of 100 µL of the supernatant was measured.

Seawater was collected from Bogliasco, Italy (44.377477, 9.070292) and filtered through a 5 µM filter to obtain the bacterial fraction. Samples were aliquoted in 30% (v/v) glycerol and stored at -80 °C for future use as inoculum. Seawater cultures were inoculated at a 1:100 ratio into fresh medium, either with 2 g/L chitin MBL or MBL with 1mM each of pyruvate, glutamate, glucose, acetate, succinate, and 1mM ammonium chloride.

### pH controlled bioreactor

A single pH-controlled bioreactor was set up in a custom-built incubator at 27°C. The reactor was a 500 mL Erlenmeyer flask with 100 mL of MBL containing 2 g/L colloidal chitin and 50 mM sodium bicarbonate instead of TES buffer. As the bicarbonate equilibrated with the environment and increased the pH, 0.2 M sulfuric acid was titrated to pH 7.8 once the culture reached pH 8.1. The acid was in a separate bottle with a pierceable rubber cap (Fisher, 15896921). A feed tube transported acid between the feed bottle and the Erlenmeyer flask using a peristaltic pump (Ismatec). A stainless steel needle (18 gauge, 6-inch stainless steel 304 syringe needle, Sigma Z102717) withdrew medium from the feed bottle. This was connected to PharMed BPT tubing (2.79 mm) through a male Luer fitting for 1/8-inch tubing (Sigma 21016). This tubing was connected to PharMed 2-stop tubing (0.25 mm, Ismatec 95723-12) using another 1/8-inch male Luer fitting and a 22 gauge, 51 mm metal hub needle (Hamilton HAM191022) inserted into the 2-stop tubing. This 2-stop tubing was placed through the peristaltic pump and connected to an additional length of 2.79 mm PharMed BPT tubing and another 18 gauge stainless steel needle. This needle dispensed acid directly into the bioreactor. The pH was continuously monitored by a controller (Milwaukee PRO pH Controller MC-122) that regulated the power supply to the peristaltic pump.

The species were inoculated into the reactor at a cell concentration of 1×10_7_ cells/mL. Once the culture reached early stationary phase (no OD_600_ increase between consecutive timepoints), the culture was centrifuged to collect biomass, then sterile filtered through a 0.2 µM membrane. EPS and the LMW fraction were collected as described below. Before analysis, 100 mM HCl was added to each fraction to remove residual bicarbonate. Since there is no organic buffer in the medium, we can attribute measured carbon to be biologically derived. Samples of EPS (dialyzed in water), the LMW fraction (cell broth filtered through a 3 kDa membrane), and biomass (cell pellet), were freeze dried, weighed, and homogenized. Elemental analysis (Molecular and Biomolecular Analysis Service MoBiAS at ETH) quantified total carbon in these fractions, reporting the final concentration in percent carbon by weight. The presence of chitin was ruled out by monitoring chitin concentrations in the EPS and biomass fractions.

### Purification of EPS

Culture supernatant was fractionated into EPS and LMW fractions using a 3 kDa cutoff filter. For large scale purification of 100mL or more, an Amicon stirred cell was used to concentrate the medium down to less than 20ml, where the concentrate was placed in a 1kDa dialysis membrane and dialyzed in water. Small volumes, centrifugal units were used, and rather than dialyzing, the concentrate was resuspended in water and further concentrated to dilute out salts. The result EPS concentrate was then freeze-dried and stored at -20°C until further use. The LMW fraction is the filtrate of the 3 kDa filter.

### Excitation Emission measurements and PARAFAC

Emission excitation matrices were acquired using a Tecan M plex plate reader, with excitation wavelengths between 250 and 450, and emission wavelengths between 300 and 600, measured at 5nm intervals. Coble peak integration, PARAFAC models, and component integration was performed using staRdom_41_ R package.

### 16S sequencing of microbial communities grown on EPS

Frozen pellets containing 1 OD-mL equivalent were used for genomic DNA extraction with the Monarch Genomic DNA Purification Kit (NEB). DNA quantity and quality was then assessed using a qubit fluorometer and a nanodrop spectrophotometer and all samples were diluted to 3.33 ng/µL. For each sample, 10 ng (3 µl) of genomic DNA was used to prepare sequencing libraries using the 16S Barcoding Kit 24 v14 kit (Oxford Nanopore) following manufacturer’s instructions with a unique barcode for each sample. After library preparation, samples were pooled and sequenced on a MinION Mk1B sequencer with a Flongle adapter using R10 flongle flow cells. Reads were filtered for size and quality using Cutadapt^42^, yielding an average of ∼30,000 filtered reads per sample. The filtered reads were then used to estimate species relative abundances using Emu^43^ with default parameters.

### Protein measurements

Proteins were quantified by adding 30 µL of sample to 300 µL of Bradford reagent (Thermo Scientific Pierce Bradford Protein Assay Kit) and quantification used standards of bovine serum albumin.

### EPS digestion using enzyme cocktails

Here we adapted a protocol based on a previous approach using microbial enzyme cocktails to degrade and characterize polysaccharides^44^. Three milliliters of EPS-containing medium (1 g/L in MBL with 1 mM ammonium chloride) were inoculated with seawater and grown at 27 °C for 72 hours until saturation. Cells were centrifuged, and the supernatant was sterile-filtered using a 0.2 µM filter. Enzymes were concentrated 10-fold using a 10 kDa Amicon centrifugal unit, with excess salts and small molecules removed by adding water and reconcentrating, repeated three times. This procedure was done in parallel for EPS substrates from each of the three degraders, yielding three sets of enzyme cocktails. The enzyme assay included 10 mM ammonium bicarbonate pH 7.8, 1 g/L EPS from one degrader, and the enzyme cocktail. The reaction was initiated by adding 50 µL of enzyme to 120 µL of the other components and was performed in parallel with a no-enzyme control to account for non-enzymatic hydrolysis of EPS. The reaction was performed with four replicates, where degradation products in three replicates were monitored in real time using LC-QTOF-MS for 24 hours. The fourth replicate was frozen immediately after the 24-hour incubation and later analyzed to acquire MS/MS spectra for compound identification.

### LC-MS measurements

The enzyme assay was initiated in the autosampler of an Agilent Infinity II 1290 multisampler and continuously sampled for 24 hours. The LC-MS method used an Agilent Poroshell PFP column (50 x 2.7 mm, 1.9 µM) at 30°C. Mobile phase A was 100% water with 0.1% formic acid (v/v), and mobile phase B was 10% acetonitrile with 0.1% formic acid (v/v). The method started with 100% phase A for 1 minute, decreased to 70% A over 30 seconds, then to 10% A over another 30 seconds, held for 30 seconds, and returned to 100% A for 3 minutes of equilibration before the next injection. An Agilent 6546 Q-TOF mass spectrometer measured product formation, with each sample injected in negative mode, followed by positive mode. Samples were acquired in high resolution mode, 50-1700 *m/z* range, 6 Hz scan speed, fragmentor 110 V, drying gas 10 L/min, and capillary voltage of 3500 V.

Peak detection and integration were performed using MZmine version 3.4.1^45^. Ion features were aligned across all samples, and their intensities were compared over time to identify ions with changing concentrations. Specifically, ions that exhibited an increase over time—defined as those with a log_2_ fold change in intensity >10 when comparing the 24-hour time point to the initial time point in one of the enzyme digests—were selected for further analysis. An inclusion list of these increasing ions was generated to acquire MS^2^ spectra for compound class identification. The same LC-MS method was employed as previously described. MS^2^ spectra were acquired using collision-induced dissociation with a collision energy of 25 volts. Data acquisition was performed with a dwell time of 50ms, a cycle time of 500ms, and a ∼1.3 *m/z* isolation window.

### Mock *Vib*1A01 EPS medium

Medium mimicking the proteins and polysaccharides present in medium containing 1g/L Vib1A01 EPS contains galactose (13 µM), Galacturonic acid (17 µM), Glucosamine (22 µM), Glucose (20 µM), and 150mg/L Bovine Serum Albumin.

### Materials and Methods NMR

Samples of 20 mg EPS have been dissolved in 150 uL D_2_O (cat. number 756822). The reference samples were 250 mM GlcNAC (Sigma) and mM chitobiose (Omicron Biochemicals cat. Number DIS-013) in 150 μl D_2_O. The solutions were transferred to 3 mm NMR tubes (Norell, S-3-HT-7) and NMR data were acquired at 303.0 K at a 600 MHz Bruker AVNEO NMR spectrometer equipped with a CP-TCI-H-C/N-D 05 Z probe head.

1D ^1^H experiments were acquired with 8192 complex data points and 32 transients. The remaining solvent signal was suppressed by applying pre-saturation or the watergate sequence with soft selective pulses^46^.

2D ^1^H-^13^C HSQC experiments were acquired with 2048 complex data points in the direct dimension, 16 transients, and 320 data points in the indirect dimension. For data acquisition and analysis, the program Topsin4.0.7 (Bruker Biospin, Inc.) was used. Data were multiplied with a squared Cosine bell window function and zero-filled to two times the original data size.

### Monosaccharide analysis of EPS

EPS of bacterial supernatants were acid hydrolysed with 1 M HCl at 100°C for 24h. Additionally, each sample was spiked with an internal standard of 15 µM 13C6-Glucose, 13C6-Galactose and 13C6-Mannose (mass 186 Da). Samples were neutralized with equimolar amounts of NaOH. Released monosaccharides (5 µL) were diluted with 20 µL of LC-MS grade water and subsequently derivatized with 75 µL of 0.1M 1-phenyl-3-methyl-5-pyrazolone (PMP) in 2:1 methanol:ddH2O with 0.4 % ammonium hydroxide, following a previously published protocols^47^. For quantification, we derivatized a serial dilution of a standard mix containing Galacturonic acid, D-Glucuronic acid, Mannuronic Acid, Guluronic Acid, Xylose, Arabinose, D-Glucosamine, Fucose, Glucose, Galactose, Mannose, N-Acetyl-D-glucosamine, N-Acetyl-D-galactosamine, N-Acetyl-D-mannosamine, Ribose, Rhamnose and D-galactosamine. Samples and standards were derivatized by incubation at 70°C for 100 minutes. After derivatization, samples were neutralized with HCl followed by 1:50 dilution in LC-MS grade water with 0.1% formic acid.

Following Xu et al.^48^ PMP-derivatives were measured on a SCIEX qTRAP5500 and an Agilent 1290 Infinity II LC system equipped with a Waters CORTECS UPLC C18 Column, 90Å, 1.6 µm, 2.1 mm X 50 mm reversed phase column with guard column. The mobile phase consisted of buffer A (10 mM NH4Formate in ddH2O, 0.1% formic acid) and buffer B (100% acetonitrile, 0.1% formic acid). PMP-derivatives were separated with an initial isocratic flow of 15% Buffer B for 2 minutes, followed by a gradient from 15 % to 20% Buffer B over 5 minutes at a constant flow rate of 0.5 ml/min and a column temperature of 50°C. The ESI source settings were 625°C, with curtain gas set to 30 (arbitrary units), collision gas to medium, ion spray voltage 5500 (arbitrary units), temperature to 625°C, Ion source Gas 1 &2 to 90 (arbitrary units). PMP-derivatives were measured by multiple reaction monitoring (MRM) in positive mode with previously optimized transitions and collision energies. Different PMP-derivatives were identified by their mass and retention in comparison to known standards. Technical variations in sample processing were normalized by the amount of internal standard in each sample. Peak areas of the 175Da fragment were used for quantification using an external standard ranging from 100 pM to 10 µM.

### MS2-based compound class annotations

For MS^2^-based annotation, fragmentation spectra were extracted using MzMine (version 4.3.0)^45^. Parameters used for mass detection were a minimal intensity of 500 for MS^1^ and 100 for MS^2^. The feature list was built using the MS^n^ tree builder with a tolerance of 20 ppm. Only features with associated MS^2^ were kept. Isotopes were searched within a 3 ppm tolerance window, features lists were aligned using a 0.1 min and 8 ppm tolerance window. Finally, the feature list was deduplicated using a 20 ppm and 0.1min window and exported for later use in SIRIUS (with a 5 ppm tolerance).

Extracted spectra were then submitted to SIRIUS^49^ v 6.0.7 and CANOPUS^38^ module for compound class annotation. All parameters used were the software default ones.

The obtained annotations were then filtered using a custom R script that combined the SIRIUS results tables together and only kept results with at least 0.5 intensity explained, an isotope score of at least 5 if present, and a most specific class probability above or equal to 0.75. The resulting table is available as **(Supplemental Dataset 3)**.

## Data availability

All mass spectral data has been deposited in the MassIVE database and will be made publicly available upon manuscript acceptance, with the password reviewer123. The kinetic data from the time course enzyme assay is MSV000096428, and the MS^2^ data for endpoint measurements is in MSV000096444.

